# A compositional reanalysis of poly(A)-selected RNA-seq reveals tank effects but no survival-associated differences

**DOI:** 10.1101/2025.04.15.648526

**Authors:** Marie Saitou, Arturo Vera Ponce De Leon, Akira Harding, Celian Diblasi, Domniki Manousi, Jun Soung Kwak, David G. Hazlerigg T, Turid Mørkøre, Lars-Gustav Snipen

## Abstract

Poly(A)-selected RNA-seq datasets are routinely generated in aquaculture research, yet the microbial information contained in unmapped reads is seldom explored due to the low abundance of nonhost transcripts and concerns about contamination. In this study, we repurposed Atlantic salmon gill RNA-seq data to assess whether meaningful microbial signals can be recovered using a contamination-aware and compositionally appropriate framework. Unmapped reads were analyzed with a custom Kraken2 database composed exclusively of complete, circularized salmon-associated bacterial genomes together with all available Atlantic salmon assemblies and the human genome. Although microbial sequences represented only a small fraction of total reads, 21 genera were detectable across samples. Genus-level profiles, Jaccard-based ordination, and ANCOM-BC analyses consistently revealed clear differences between tanks, whereas no associations were observed for sex or survival status. Three species exhibited significant tank-specific effects, indicating that environmental factors contributed the strongest detectable structure in the data. The limited microbial diversity recovered here reflects the expected constraints of poly(A)-enriched libraries, yet the results demonstrate that unmapped reads from host-derived RNA-seq can still provide informative environmental signatures when analyzed with curated reference databases and compositional statistical approaches. This strategy offers a practical means to extract exploratory microbiome information from existing transcriptomic datasets.

## Introduction

Microbial dynamics in aquaculture systems strongly influence fish health, environmental stability, and pathogen transmission (1). Although high-throughput sequencing has greatly advanced the characterization of aquatic microbiomes, most studies rely on dedicated 16S rRNA gene or metagenomic approaches, which are not always available for large production facilities or retrospective analyses (2). In contrast, host-focused RNA-seq datasets are routinely generated in aquaculture research and breeding programs yet the microbial information contained in unmapped reads is typically overlooked.

Poly(A)-selected RNA-seq is not designed for microbial profiling because bacterial transcripts generally lack polyadenylation and because microbial RNA represents only a small fraction of total RNA in host tissues (2, 3). Even so, unmapped reads in such datasets may retain detectable microbial signatures when analyzed with contamination-aware reference databases and compositional statistical methods. Recovery of such signals is particularly relevant in aquaculture, where operational and environmental conditions, including water management and tank configuration, can generate distinct microbial environments that influence fish health and pathogen exposure (4, 5).

Given that dedicated microbial sequencing is not always feasible, especially in commercial or high-throughput production settings, repurposing existing host-derived RNA-seq datasets offers a practical and cost-effective means to explore microbial information (6). Such an approach may help identify environmental signatures without requiring additional sampling.

Here, we repurposed poly(A)-selected RNA-seq data from Atlantic salmon gill tissue to evaluate the extent to which microbial information can be recovered and whether such information reflects biological or environmental structure. Using a contamination-aware Kraken2 database and compositional analyses, we characterized the microbial taxa detectable in unmapped reads and assessed their variation across sex, survival status, and tank identity. While microbial signals were limited in quantity, they were sufficient to reveal consistent and statistically supported differences between tanks, whereas no associations were observed for sex or survival. These findings illustrate both the constraints and the utility of host-derived poly(A)-selected RNA-seq data for exploratory microbial analyses in aquaculture systems (7–9).

## Materials and Methods

### Metatranscriptome dataset

The metatranscriptome dataset analyzed in this study was derived from Atlantic salmon gill samples collected in June 2021, one week prior to seawater transfer, at which time the fish showed no visible signs of disease. These fish originated from 109 full-sib families of the Mowi (Mowi Genetics AS) strain, which had been generated through selective breeding in November 2019. The metatranscriptome dataset is publicly available at the European Nucleotide Archive (ENA) under project number PRJEB47441. This study did not involve direct handling of fish; instead, it retrospectively analyzed transcriptomic data to investigate microbial dynamics prior to disease onset. A comprehensive description of the fish population is provided in (10) while details on metatranscriptome data generation are outlined in (11). The original experiment was performed according to EU regulations concerning the protection of experimental animals (Directive 2010/63/EU). Appropriate measures were taken to minimize pain and discomfort. The experiment was approved by the Norwegian Food and Safety Authority (FOTS id. number 25658) (10).

### Fish Rearing and Sampling Timeline

Fish were initially maintained under continuous light (LL) from hatching until they reached approximately 50g. They were then exposed to a 6-week 12:12 photoperiod (12 hours light, 12 hours dark) before returning to LL for 8 weeks, reaching 100g, the standard size for seawater transfer.

In early February 2021, fish were vaccinated with Alpha Ject micro 6 (PHARMAQ®) against *Aeromonas salonicida, Listonella anguillarum O1* and *O2a, Vibrio salmonicida, Moritella viscosa, and IPN virus Sp*. In June 2021, one week before seawater transfer, gill biopsies were collected for RNA sequencing at a time when fish showed no visible disease symptoms.

On June 23, 2021, fish were transported to the ARST seawater facility. Approximately eight weeks after transfer, signs of winter ulcers began to appear, and mortality increased throughout the autumn. Despite the removal of affected individuals during trait recordings in early November, the prevalence of winter ulcers persisted into the winter. At the end of November, phenotypic data were recorded, marking the conclusion of the experimental phase.

### Data Selection

To allow for a balanced investigation of microbial communities in the context of fish health, sample selection was based on multiple biological and experimental constraints. The dataset included 15 surviving fish, alive at five months post-transfer, and 13 non-surviving fish, which died within 40 days post-transfer (**Table 1**). Sex was evenly distributed, with 13 males and 15 females, to account for potential sex-specific microbial differences. Fish were reared in two separate tanks, C1 and C2, allowing the assessment of environmental effects. The died group consisted exclusively of fish with visible wounds, ensuring a consistent disease phenotype, while the survived group included only wound-free individuals, providing a clear contrast (**Figure 1**). While the selection aimed for a balanced representation across conditions, final sample availability was constrained by the accessibility of metatranscriptome data and the biological conditions of the cohort.

**Table 1.** Most transcriptionally represented microbial taxa in Atlantic salmon gill metatranscriptomes.

| taxon | lfc_TankC2 | se_TankC2 | q_TankC2 |
| --- | --- | --- | --- |
| *Brucella rhizosphaerae* | -1.306 | 0.322 | 0.00713 |
| *Pseudomonas qingdaonensis* | -1.321 | 0.322 | 0.00713 |
| *Pseudomonas iridis* | -1.225 | 0.319 | 0.00801 |

Species exhibiting significant differential abundance between Tanks C1 and C2 according to ANCOM-BC2. Taxa were considered significant when the differential abundance indicator for the tank contrast was TRUE and the FDR-adjusted q-value < 0.05. Negative log-fold change (lfc) values indicate higher abundance in Tank C1.

**Figure 1.**
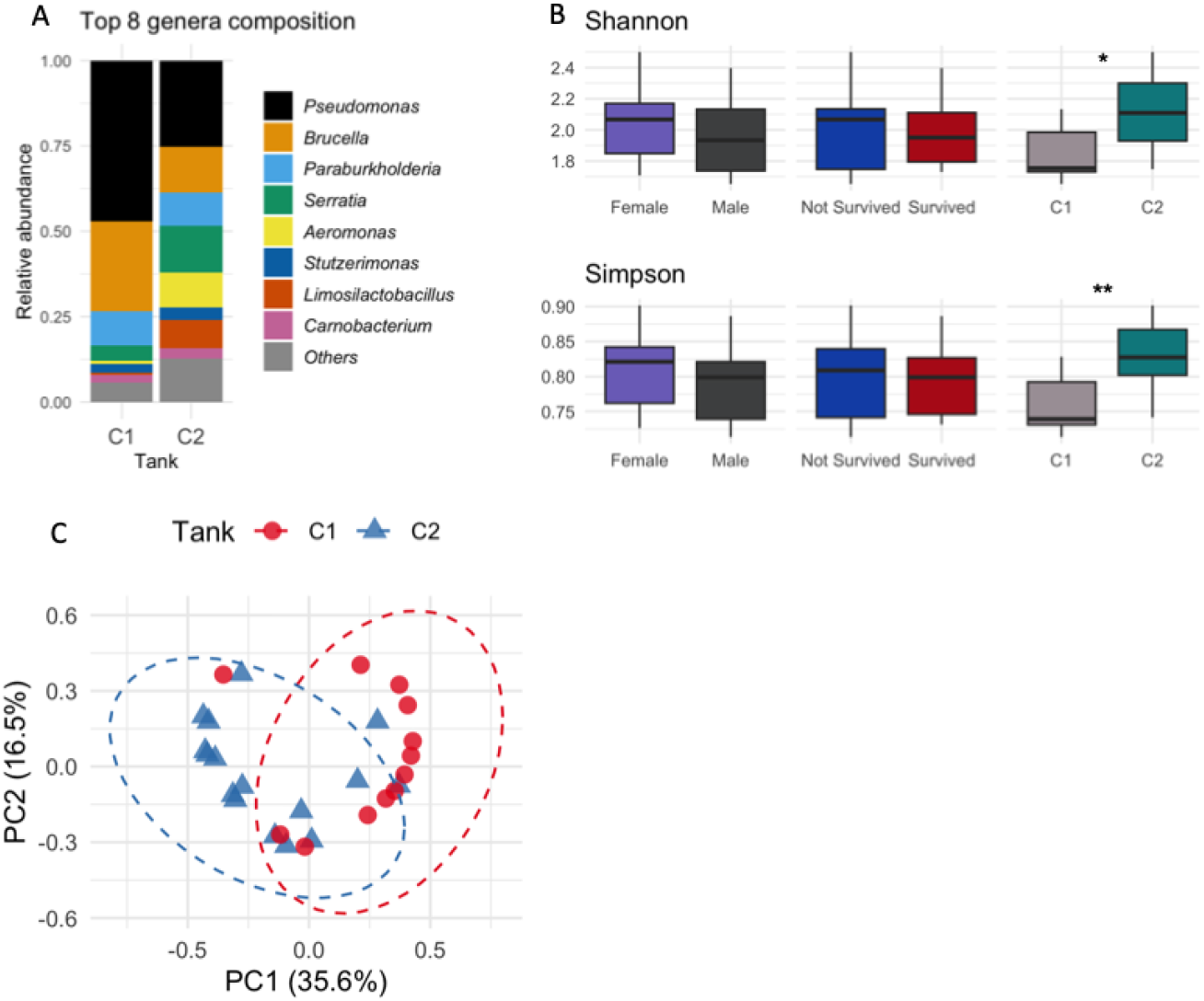
Variation in detectable microbial communities across conditions. **A. Genus-Level Microbial Composition in Tanks C1 and C2** Species-level Kraken2 counts were aggregated to the genus level after removing reads assigned to *Salmo salar* and *Homo sapiens*. The eight genera with the highest total counts across both tanks were retained individually, and all remaining genera were grouped as “Others.” Bars represent relative abundances within each tank. **B. Shannon and Simpson alpha diversity per group**. Shannon and Simpson diversity indices calculated from species-level microbial counts. Diversity values were compared across sex, survival status, and tank using Wilcoxon rank-sum tests with Benjamini–Hochberg correction. An asterisk (*) indicates q < 0.05, and a double asterisk (**) indicates q < 0.01. **C. Principal coordinates analysis (PCoA) of microbial communities based on Jaccard dissimilarity**. Each point represents one sample and is colored and shaped by tank identity. Ellipses indicate 95% confidence intervals. Samples separate according to tank, indicating that detectable microbial composition differed between tanks.

### Metatranscriptome data preparation

The RNA-seq data used in this study are publicly available in FastQ format from the European Nucleotide Archive (ENA; project PRJEB47441). The dataset consists of gill metatranscriptome from Atlantic salmon, with sequencing performed using a paired-end (PE) approach (2×150 bp) generating approximately 20 million reads per sample. Quality assessment of raw reads was conducted using FastQC (v0.12.0). Adapter sequences, low-quality bases, and short reads were removed using fastp (12)(v0.23.4) with default parameters. Filtered reads were then mapped to the Atlantic salmon reference genome (13) (Ssal_v3.1, GCF_905237065.1) using bwa-mem2 (v2.2.1). Reads that failed to align with the host genome were extracted and retained for microbiome analysis.

The RNA-seq libraries were prepared using a poly(A) selection method to enrich for mRNA, rather than actively depleting ribosomal RNA (rRNA). While poly(A) selection primarily targets eukaryotic mRNA, certain bacteria and some prokaryotic organisms possess polyadenylated mRNA at low frequencies, which may result in the retention of microbial transcripts. Additionally, host-derived poly(A) transcripts from Atlantic salmon may be present at a consistent proportion across samples.

For preprocessing, adapter trimming and quality filtering were performed using fastp (v0.23.4), followed by host genome mapping against the Atlantic salmon reference genome (Ssal_v3.1, GCF_905237065.1). Unmapped reads were retained for microbiome analysis, under the assumption that they predominantly represent microbial sequences. However, given the poly(A) selection strategy, it is possible that a proportion of these unmapped reads include artifacts from host transcripts or residual polyadenylated bacterial transcripts. Although this potential artifact is present, it is expected to be uniformly distributed across all samples rather than selectively biased toward a specific group (e.g., survived vs. died fish). Therefore, while this factor must be considered when interpreting metatranscriptomic results, the relative comparisons between survival groups remain valid, as any systematic bias introduced by poly(A) selection would be consistent across conditions.

### Construction of a custom Kraken2 reference database

A custom Kraken2 database was constructed to ensure contamination-free taxonomic classification and to minimize false-positive assignments arising from incomplete or contaminated reference genomes. The database consisted exclusively of: (1) bacterial genomes included in the Salmon Microbiome Genome Atlas (SMGA(14)); (2) the Atlantic salmon genome Ssal_v3.1 (Ref Seq assembly GCF_905237065.1); and (3) the complete human genome assembly GRCh38 (Ref Seq assembly GCF_000001405.26). All bacterial reference genomes were retrieved from https://ns9864k.web.sigma2.no/TheMEMOLab/projects/SMGA_v1.0/ and only assemblies annotated as “complete genome (higher than 75 % complete and less than 5 % contamination according to ChekM2 scores (15))” were included for the bacterial component to reduce the likelihood of cross-species contamination. The database was built in October 2025, following the standard Kraken2 build procedure. The final database contained 211 bacterial genomes, the *S. salar* genome, and the human genome. Kraken2 (v2.1.4) was used for taxonomic assignment of unmapped reads (16).

### Taxonomic assignment of unmapped reads

Unmapped reads were classified using Kraken2. For each sample, species-level counts were extracted from the Kraken2 report files by retaining entries assigned at rank “S” (species). Counts assigned to *Salmo salar* and *Homo sapiens* were removed prior to analysis. The resulting species-by-sample table was assembled by merging all Kraken2 report outputs. When species were absent from a given sample, counts were recorded as zero.

### Genus-level aggregation

Species names were parsed to extract the genus (the first element of the species name). Genus-level counts were obtained by summing species-level counts within each genus. For compositional summaries, counts were further aggregated within each tank and converted to relative abundances. To visualize dominant taxa, the eight genera with the highest total abundance across both tanks were retained as individual categories, with all remaining genera grouped as “Others.”

### Phyloseq object construction

A *phyloseq* object was created using sample metadata and the species-level count matrix. All downstream diversity and differential abundance analyses were performed within this framework.

### Differential abundance analysis

Differential abundance testing was conducted using ANCOM-BC2(17). The model included Tank, Sex, and Survival status as fixed effects. Benjamini–Hochberg correction was applied to control the false discovery rate. Taxa were considered statistically significant when the q-value was less than 0.05 and the differential abundance indicator for the corresponding coefficient was TRUE. Significant species associated with tank identity were extracted for reporting in Table 1.

### Alpha diversity

Shannon and Simpson diversity indices were computed from species-level counts after removal of host and human reads. Diversity values were compared across Tank, Sex, and Survival categories using pairwise Wilcoxon rank-sum tests within each index, followed by Benjamini–Hochberg adjustment.

### Beta diversity

Presence–absence community dissimilarity was quantified using Jaccard distances calculated from the species-level count table. Classical multidimensional scaling (principal coordinates analysis) was applied to the Jaccard distance matrix. The first two principal coordinates were plotted to visualize community structure by tank identity, sex, and survival status.

### Software

All analyses were performed in R using the packages *tidyverse, phyloseq, ANCOMBC, vegan*, and *rstatix*(18–22). Scripts used for all processing steps, including taxonomic classification and statistical analysis, are provided as supplementary material to ensure full reproducibility.

We used ChatGPT (version 5.2, OpenAI) to assist with code refactoring, language editing, and proofreading. All scientific interpretations, analyses, and conclusions are the responsibility of the authors.

The scripts are available in the Figshare repository: https://doi.org/10.6084/m9.figshare.29482994

## Results

We obtained an average of 537,995 ± 289,590 unmapped reads from a total of 11,328,436 ± 1,190,386 reads from the 28 samples. To characterize the microbial taxa detectable in the poly(A)-selected RNA-seq data, species-level Kraken2 assignments were aggregated to the genus level after exclusion of reads assigned to Salmo salar and Homo sapiens. Across all samples, 21 genera were observed. When genera were ranked by their total counts across both tanks, the most frequently detected genera included *Pseudomonas, Brucella, Paeniglutamicibacter, Micavibrio, Sphingomonas, Sphingobium, Williamsia, and Kocuria*. These eight genera were retained as individual categories, and all remaining genera were grouped as “Others.” The resulting aggregated genus-level relative abundances for Tanks C1 and C2 are shown in **Figure 1**, illustrating the proportional contribution of these genera within each tank. To summarize within-sample diversity, Shannon and Simpson indices were calculated from the specieslevel microbial count table after removing reads assigned to *Salmo salar* and *Homo sapiens*. Diversity values were compared across Sex, Survival status, and Tank categories. No differences were observed for either Shannon or Simpson diversity between Sex groups (female versus male) or between Survival groups (not survived versus survived), with Benjamini–Hochberg–adjusted P-values exceeding 0.33 for all comparisons. In contrast, both Shannon and Simpson indices differed between Tank C1 and Tank C2, with adjusted P-values of 0.012 for Shannon and 0.0049 for Simpson (**Figure 1B**). To examine sample-to-sample differences based on taxonomic presence or absence, Jaccard dissimilarity was calculated from the species-level read assignments. The first two principal coordinates are presented in **Figure 1C**. Samples from each tank are displayed according to their tank identity, allowing the spatial arrangement of samples to be viewed in ordination space. This representation indicates how samples relate to one another based on shared and non-shared microbial species.

To identify species exhibiting statistically detectable differences across sample variables, ANCOM-BC2 was applied to the species-level count table with Tank, Sex, and Survival status included as covariates. Three species showed a significant coefficient for the Tank contrast after Benjamini–Hochberg correction: Brucella rhizosphaerae (lfc = −1.306, SE = 0.322, q = 0.00713), Pseudomonas qingdaonensis (lfc = −1.321, SE = 0.322, q = 0.00713), and Pseudomonas iridis (lfc = −1.225, SE = 0.319, q = 0.00801). No species exhibited significant associations with Sex or Survival status. These taxa are listed in **Table 1**. The species identified as significant contribute to genera that appear within the genus-level profiles in **Figure 1**, although genus-level aggregation does not yield a direct one-to-one correspondence between the two representations.

In sum, C1 contained higher relative abundances of several dominant genera, whereas C2 showed a more evenly distributed genus composition. Consistent with this, alpha diversity values were higher in C2 than in C1. ANCOM-BC2 identified three species belonging to the genera observed to be enriched in C1, further reflecting the concentration of specific taxa within that tank.

## Discussion

In this study, we examined microbial signatures detectable in poly(A)-selected RNA-seq data from Atlantic salmon by reanalyzing unmapped reads using a contamination-aware taxonomic approach. After removing reads assigned to the host and human genomes, only a modest number of microbial reads remained, consistent with the expectation that poly(A) enrichment does not target prokaryotic transcripts. While poly(A) selection bias is expected to be systematic across samples, it may still differentially affect taxa depending on the degree of transcript polyadenylation (23, 24). Thus, despite inherent biases, meaningful comparative analyses remain possible within the detectable subset of microbial transcripts. Despite these constraints, the detectable microbial features revealed clear differences between tanks, whereas neither sex nor survival status showed any evidence of association.

The genus-level community profiles and Jaccard-based ordination both indicated that samples clustered according to tank identity. This pattern was further supported by differential abundance testing using ANCOM-BC2, which identified three species with significant tank-associated effects. No species were associated with sex or survival status. These results suggest that, within the limits of a poly(A)-selected dataset, environmental factors related to tank conditions exert a stronger influence on detectable microbial composition than fish-level biological traits (25).

The taxonomic composition observed here differed in part from profiles typically reported in 16S rRNA gene sequencing or shotgun metagenomic studies of Norwegian aquaculture environments. While some genera detected in this study, such as *Pseudomonas*, are also commonly reported components of marine and aquaculture-associated microbiota, their relative abundances differed from those observed in previous studies(26). In contrast, other taxa identified here, including *Brucella*, are not typically reported as abundant members of aquaculture microbiomes and are rarely detected in comparable surveys. These discrepancies likely reflect methodological differences, including the use of poly(A)-selected RNA-seq data and stringent contamination-aware filtering, which together constrain the subset of microbial transcripts that can be recovered. Combined with the use of a contamination-controlled database, these factors likely restrict detectable diversity and shift relative abundances away from patterns observed with less biased methods. Accordingly, the results presented here should be interpreted as reflecting the subset of microbial signals that remain after host-dominated transcriptional data have been filtered rather than as comprehensive estimates of microbial community structure. The importance of using contamination-controlled reference databases has been underscored in other RNA-seq–based microbiome studies. In particular, incomplete or contaminated bacterial genomes can generate extensive false-positive assignments in human tumor RNA-seq data (27), and that restricting the database to validated complete genomes markedly reduces such artefacts. This methodological consideration is directly relevant here and provides context for the limited diversity and shifted relative abundances observed in our dataset

Overall, our findings illustrate that microbial information present in poly(A)-selected RNA-seq data, although limited, can still capture robust environmental signatures when analyzed with appropriate contamination-aware and compositional methods. Microbiome count data are inherently compositional, such that analyses based on unadjusted relative abundances can lead to misleading inferences (28). This consideration further supports the use of ANCOM-BC2 in the present reanalysis and contextualizes the tank-associated differences detected here. The tank-specific differences detected here demonstrate that unmapped reads can provide biologically meaningful insights into environmental influences on salmon-associated microbial communities. Future work comparing poly(A)-selected datasets with rRNA-depleted metatranscriptomes or metagenomics may help to clarify how methodological choices shape the detectable microbiome and to establish best practices for repurposing host-focused RNA-seq data for microbial investigations.

## Author contributions (CRediT)

Akira Harding: Conceptualization, Validation, Formal analysis, Investigation

Celian Diblasi: Conceptualization, Methodology, Investigation, Writing – Original Draft

Domniki Manousi: Data curation

Jun Soung Kwak: Conceptualization, Methodology

David Hazlerigg: Resources, Funding acquisition

Turid Mørkøre: Resources, Visualization

Arturo Vera Ponce De Leon: Conceptualization, Methodology, Supervision

Lars-Gustav Snipen: Methodology, Supervision

Marie Saitou: Conceptualization, Investigation, Writing – Review & Editing, Supervision

## Data Availability Statement

The metatranscriptomic sequence data analyzed in this study are publicly available at the European Nucleotide Archive (ENA) under project number PRJEB47441.

The scripts and annotation files used for taxonomic and functional profiling are available in the Figshare repository: https://doi.org/10.6084/m9.figshare.29482994

No new sequencing data were generated as part of this study.

## Acknowledgements

The authors thank the Galaxy.no and Orion high-performance computing infrastructures for providing computational resources. We are also grateful to Côme Morel (CIRAD-BIOS, UMR ASTRE) for his valuable advice on alpha and beta diversity analyses.

## Tables and Figures

